# Variation in social feeding behavior and interactions among *Caenorhabditis* nematodes

**DOI:** 10.1101/2025.06.26.661336

**Authors:** D Haskell, J Palo, RFH Eugene, CRL Large, MP Hart

**Affiliations:** Department of Genetics, Perelman School of Medicine, University of Pennsylvania

## Abstract

The ability to respond to complex stimuli and environmental cues is essential for organisms to survive and reproduce. Responding to a wide range of stimuli requires a neuronal network that can integrate cues and execute behavioral responses. Evolution of behaviors occurs ubiquitously in most established ecological niches, even among closely related species. To uncover the genetic and neuronal drivers of evolving behaviors, we have taken advantage of the large and relatively ancient divergence in the *Caenorhabditis* genus to ask how different *Caenorhabditis* nematodes respond to environmental stimuli and whether behavioral traits are shared or distinct. Here, we assayed foraging behaviors of twelve members of the *Caenorhabditis* clade, including members of both the *elegans* and *japonica* supergroup, and the basal taxon *C. monodelphis*. For each species, we analyzed social feeding and bordering behaviors, which are well characterized in *C. elegans.* These behaviors are the functional readout of complex sensory integration of multiple sensory cues including pheromones, touch, O_2_/CO_2_ concentration, and attractive and noxious stimuli. We hypothesized that the evolutionary divergence between species would correlate to divergence in these behaviors. We observed a wide variation in social aggregate feeding and bordering behaviors of hermaphrodite and female animals, but the variation did not correlate with evolutionary relatedness of the species. Addition of male animals with female or hermaphrodite animals of the same species increased aggregation behavior of subset of species, but not others. Combination of a second species with *C. elegans* significantly reduced aggregate feeding behavior of *C. elegans*, but not the other species. Intraspecies and interspecies interactions therefore modifies behavioral paradigms. Overall, we find that foraging and social feeding behaviors vary widely across *Caenorhabditis* species, likely due to species-specific responses and integration of environmental and context sensory cues. In general, the clade represents a compelling model to dissect evolution of behavior across diverse environments and a large timescale.

## Introduction

Behavior is how organisms interact with their environment and respond to both internal and external stimuli. Simple forms of life, especially those that represent unicellular phyla, rely heavily on the detection of chemical and tactile stimuli through direct contact^1^. In contrast, broad sensory specialization is heavily emphasized in Eukaryotes, which range in the complexity of their specialization (including the evolution of differentiated and highly specialized neuronal tissues) but also in the manner and breadth of their behavioral responses to the environment^2,3^. The evolution of complex behavior can be loosely correlated with the evolution of multicellularity and complex tissues^4^. Likewise, tissue differentiation also gave rise to neural tissues, which in their simplest form, link networks of sensory cells together forming a basic neural circuit. In more complex Eukaryotes, more sophisticated neural networks, utilizing multiple sensory and neuronal cell types form the basis of complex organs such as neural ganglion, nerve tracts, and the brain^5,6^

Changes in behavioral traits occurs regularly in most eukaryotic organisms, especially between related species^7^. Logically, rapid shifts in behavior should be tightly coupled with the congruent alteration of their physiological and genetic drivers as highlighted by several studies in *Drosophila* and related species^8,9^. Small changes in gene regulatory networks, modest alterations in neural circuits, or mutations influencing receptor/ligand interactions are all sufficient to alter behavioral paradigms in an often-unpredictable manner. Without the ability to identify stepwise drivers of shifting behavioral paradigms it can be challenging to understand how selective pressures lead to modification of underlying genetic and molecular factors, which ultimately dictate the neuronal activity underlying complex responses to stimuli.

Compounding the study of behavioral evolution is the complex nature of behavior itself. To truly differentiate between behavioral paradigms requires dissection at multiple levels of organismal complexity, starting with changes to the genome and epigenome, and culminating in intra-circuit alterations leading to modifications in the complex network of integrated and often co-dependent behaviors. While comprehensive dissection of genetic and molecular drivers of altered or novel behaviors is rarely achieved, we can utilize existing genetic information in clades such as *Caenorhabditis* to inform research to begin circuit level descriptions.

Amongst common lab models, the nematode *Caenorhabditis elegans* is a particularly powerful tool for studying neuronal circuits and behaviors. In addition to the many genetic tools, a fully mapped neuronal connectome (including chemical, electrical, and peptide signaling), provides a powerful model to link stereotyped behaviors with alterations of neuronal activity at both a single cell and circuit level. In addition, the *Caenorhabditis* clade of nematodes represents a unique opportunity to study the evolution of behavior in a diverse and evolutionarily ancient lineage **(Figure 1)**^10^. Nematodes of this clade are found worldwide, with populations found in nearly every ecological niche probed for their presence^11^. In the wild, members of the *Caenorhabditis* nematodes persist in a free-living state generally colonizing bacteria-rich substrates, such as rotting fruit, allowing them to utilize the local microbiota as a food source. Considering their global dispersion, individual members of the clade experience significant variation in local environment, including differences in temperature, humidity, concentration of oxygen, and dominant microbiota^12^. Notably, these species are rarely isolated from homogenous populations; for example *elegans* is frequently co-isolated with *briggsae* and *remanei* from the same substrate. Altogether, the *Caenorhabditis* clade presents a compelling model through which to query the evolution of behavior across a large evolutionary timescale and a wide range of environments.

**Figure 1.**
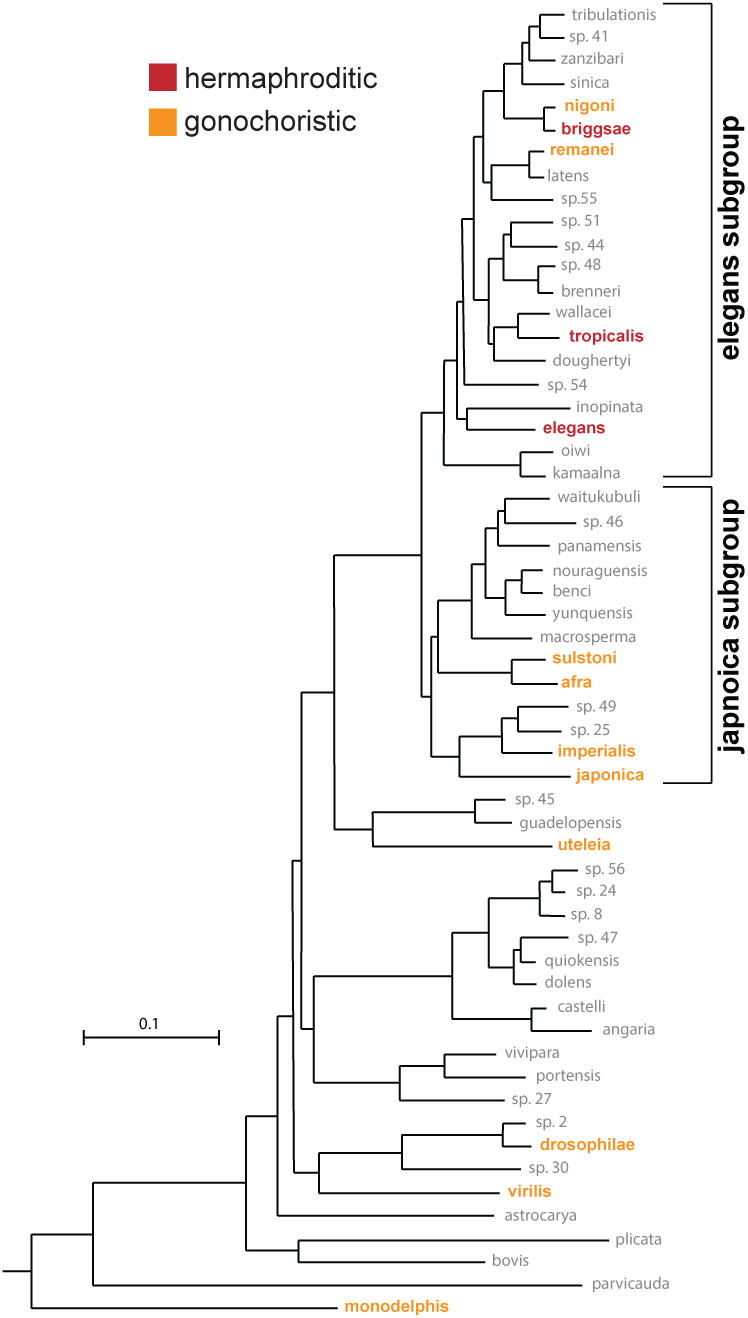
The *Caenorhabditis* clade of nematodes is an evolutionary and diverse lineage. **(A)** Phylogenetic tree of *Caenorhabditis* clade showing the two large subgroups (*elegans* and *japonica*) and the basal taxon *monodelphis*. Species included in behavioral assays are highlighted and color-coded based on reproductive strategy (red = hermaphroditic or orange = gonochoristic). Figure was adapted from Stevens et al., 2020^10^.

Due to high levels of variation among local environments, we chose to first assay foraging behaviors, which we reasoned would be ubiquitously present among the clade given likely similar or shared feeding strategies. One of the most well-characterized feeding behaviors is termed social feeding and is defined by social interactions that occur between individuals feeding in the same local environment^13^. Although not well defined, one can speculate that this behavior has costs and benefits for individual animals and for populations. Animals may be attracted to high density food that is resource rich for obvious nutrient seeking reasons, but animals populating a food/bacteria rich location can also result in spreading of the food source (bacteria) to surrounding areas. Further, group feeding could facilitate short-distance signaling between animals (perhaps through pheromones) and/or increase opportunities for mating, but it seems clear that this behavior is environment and context dependent^14,15^. Early surveys of wild *elegans* isolates showed a surprisingly high level of variability in social feeding behaviors. Contrary to the widely used N2 Bristol lab strain, which displays solitary feeding, many wild isolates from a range of environments show higher levels of social feeding^13^. de Bono & Bargmann defined the solitary feeding behavior of N2 Bristol strain to a gain of function missense mutation in the *npr-1* gene (ortholog of neuropeptide Y receptor)^13^. This results in an amino acid change in the third intracellular loop of the NPR-1 G-protein coupled receptor. Loss of function mutations in the *npr-1* gene result in social feeding in *elegans*^13^. For the purpose of this study we utilize the *npr-1(ad609)* loss of function mutant to induce social feeding in the N2 lab control strain, which represents a reliable and robust positive control for *elegans* social feeding behaviors observed in wild isolate *elegans* strains^16^.

Subsequent work further elucidated the role of NPR-1 and other neuronal drivers of solitary and social feeding, including that *elegans* social feeding behavior is driven primarily through the integration of environmental cues including pheromones, O_2_/CO_2_ concentrations, and perception of touch and noxious stimuli ^14,17–20^. Sensory neurons including ADL, ASH, ASK, ADE and AWB form a circuit to detect the local sensory environment which is then integrated primarily through the interneuron RMG to alter locomotion paradigms^17,18^. Genetic dissection of this sensory circuit has uncovered distinct roles for the neuropeptide receptor NPR-1^13,16^ and gap junction connections between involved sensory neurons and the RMG interneuron^17,21^. More recent work has implicated glutamate signaling from ADL and ASH sensory neurons, where conserved synaptic adhesion molecules (NRX-1 and NLG-1) contribute to social feeding, and *nrx-1* impacts architecture and function of ASH chemical synaptic connections^16^. Therefore, social feeding behavior relies on co-dependent signaling of neuropeptides, and electrical and chemical synapses, which together fine-tune the balance between solitary and social feeding behaviors^16^. Given the complexity of this behavioral circuit, we asked whether social feeding behaviors are conserved throughout the *Caenorhabditis* clade of nematodes.

In this study, we compared social feeding behaviors of twelve *Caenorhabditis* species to multiple strains of *elegans* (**Figure 1**). This clade of nematodes is not only relatively ancient, but also has significant divergence over evolutionary time, with an estimated genomic divergence of > 20 million years between *elegans* and the four most-closely related species^22^. Two large subgroups, *elegans* and *japonica*, have emerged based on genomic data providing multiple lineages to interrogate^10^. *monodelphis*, representing the basal taxon of the clade, is estimated to have diverged over 100 million years ago, an evolutionary timescale far beyond that of the split between mice and humans ^23^. Lastly, the true strength of this model lies in the opportunity to study “wild” populations of nematodes (representing both hermaphroditic and gonochoristic species) in a controlled lab setting applicable to medium-throughput behavioral assays. When considering the evolution of behavior within the *Caenorhabditis* clade we set out to compare differences amongst the species, providing a base on which future work investigating differences in sensory cues, neuronal function, and circuitry can be launched.

Using the twelve members of the *Caenorhabditis* clade we found significant variation in social feeding behaviors of hermaphrodites and females amongst the different species, with all but two showing reduced levels of aggregate feeding compared to the *elegans* social feeding control strain, and half of the species significantly increasing social feeding than the *elegans* solitary control strain. Furthermore, by testing more ecologically relevant populations in our assays we identified the impact of intraspecies and interspecies interactions. Mixing males with females (or hermaphrodites) of individual species altered aggregate feeding behavior in two species when compared to homogeneous female (or hermaphrodite) populations. Combining a second species with social feeding *elegans* was sufficient to significantly reduce aggregation feeding of *elegans*. This work represents the first characterization of foraging and social feeding behaviors across the *Caenorhabditis* clade of nematodes and lays the groundwork for the dissection of ecological, cellular and circuit level mechanisms of conserved behaviors.

## Methods

### Strains Maintenance

Animals were maintained under normal growth conditions (∼20° C) on normal NGM media^24^. Plates were seeded with *E. coli* OP50 to provide food. Strains utilized in this study are listed in **Table 1**. *npr-1*(*ad609*) and *nrx-1*(*wy778*) mutations were confirmed in *elegans* strains and were used as positive controls. All other species were obtained from the CGC and maintained by chunking.

**Table 1.**
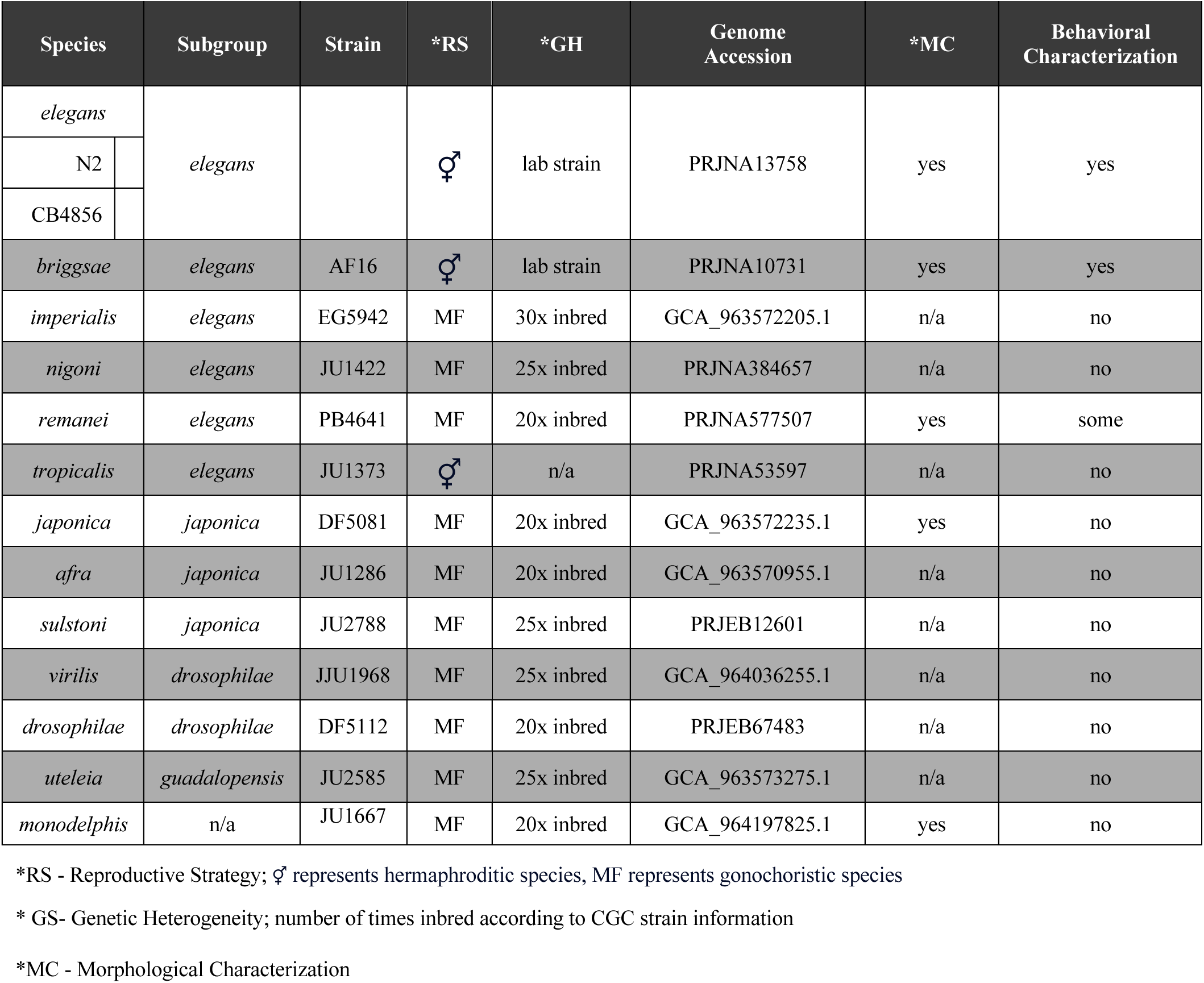
Reference table for *Caenorhabditis* species information.

### Social feeding behavior assay

Standard 6-well culture plates were filled with 6 mLs of standard NGM media and then seeded with 75 μL of *E. coli* OP50 and allowed to dry overnight. For the standard assay, 50 animals were picked onto clean plates and then moved onto the seeded wells, transferring as little bacteria as possible to prevent buildup. For standard social feeding and bordering assays (**Figure 2B&C**) the 50 animals were either female or hermaphrodites depending on the species. For the mixed sex assays, we combined 25 males with 25 females/hermaphrodites (depending on species). Regardless of the composition of the assay, worms were picked during and placed on the assay during their fourth larval stage. Female or hermaphrodite animals were staged based on the presence of half-moon shape characteristic of a L4 developing vulva. Males were staged based on relative size, germline features, and morphological development of the tail. After placing the animals on the 6-well plate, a small amount of Tween20 was used to thinly coat the lid to prevent condensation forming during imaging. The plates are imaged using the WormWatcher automated imaging platform (Tau Scientific) for a total of 20 hours with a cluster of 10 images (over 1 minute timecourse) being taken every hour. To quantify social feeding behavior each well was manually scored from blinded images, where a worm was considered aggregating if it was contacting two or more other worms^13,16^. Bordering was assayed by calculating the percentage of solitary animals in direct contact with the border region of the bacterial lawn (with the border defined as approximately half a body length in width). Data shown is from the 15-hour mark, which placed the animals of all species in the middle of Day 1 of adulthood.

**Figure 2.**
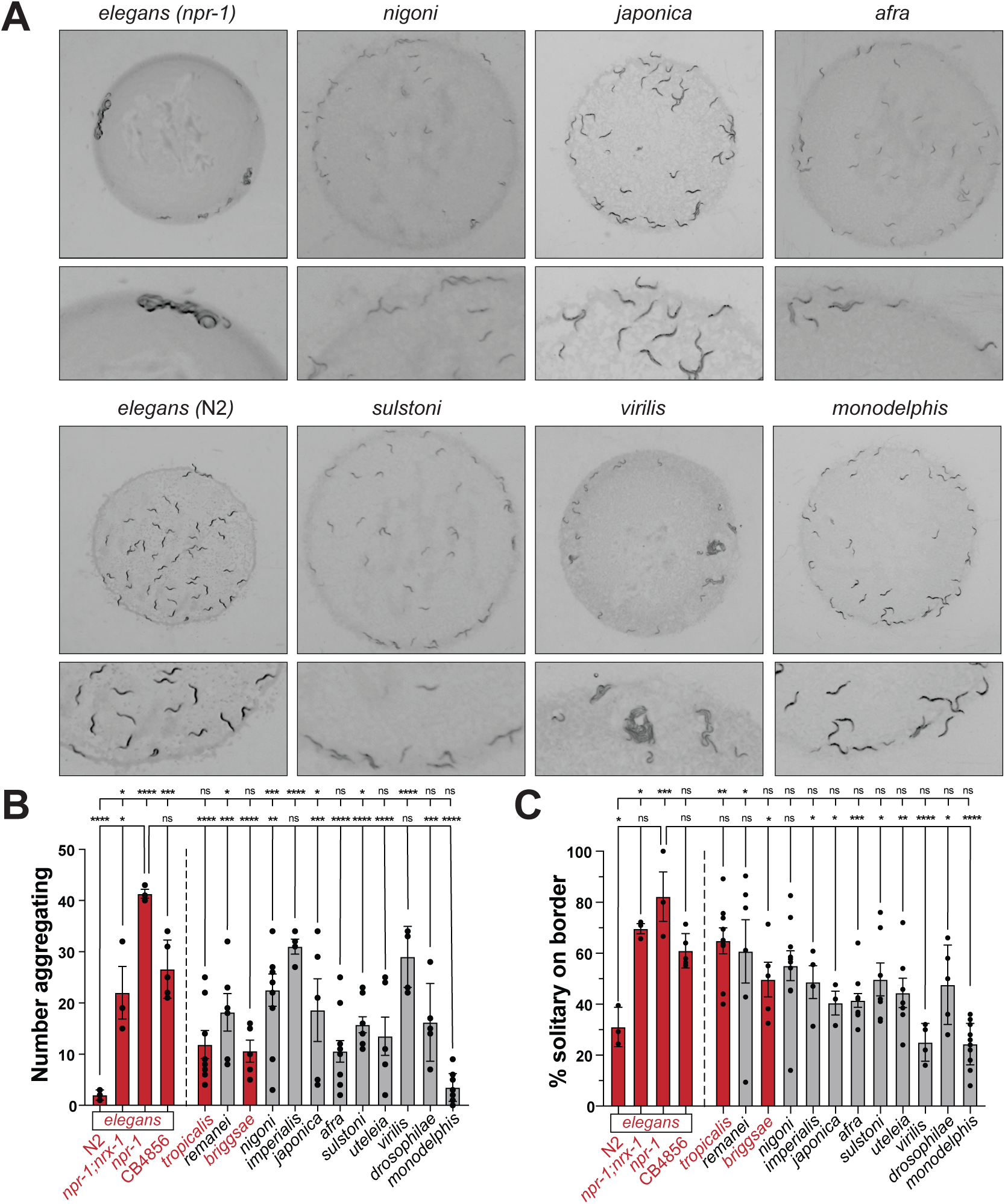
Aggregation behavior is diverse across *Caenorhabditis* clade. **(A)** Representative images of select *Caenorhabditis* species aggregation feeding and bordering behaviors. **(B)** Quantification of number of aggregating animals in 13 members of the *Caenorhabditis* clade. Species are ordered based on the provided phylogeny (and for subsequent figures). Significant variation is present among members of the clade, and also within individual species. The 4 *elegans* groups assayed represent the negative control (N2 ‘Bristol’), positive control *npr-1(ad609)*, modulated positive control *npr-1(ad609*);*nrx-1*(*wy778*), and wild isolate CB4856 ‘Hawaiian’, respectively. Bar graphs represent the mean values across biological replicates (each replicate indicated by a dot). Error bars show stand error of mean. One-way ANOVA with Tukey’s post-hoc test was used for statistical comparisons, with p-values indicated in standard format (p=0.05(*), p=0.01(**), p=0.001(***), p= <0.001 (****)). Hermaphroditic species are labeled in red, while gonochoristic species are labeled in black. **(C)** Quantification of percent solitary animals on food border in 13 members of *Caenorhabditis* clade. Significant variation is present among members of the clade, and also within individual species. Graphical representations and statistical analysis are consistent with previous panel.

### Mixed species assays

In order to assay the effect of mixed species, we first generated a fluorescently tagged *elegans* strain to differentiate them from the other species while imaging behavior. *npr-1(ad609) elegans* were injected using standard technique with plasmid *myo-2p*::*mScarlet* in order to label pharyngeal muscle (25 ng/μL). Strong expression of the transgene allows visualization of a single worm when in an aggregate. To perform the assay a standard 5cm NGM plates was seeded with 25 μl of *E. coli* OP50 and allowed to dry. To mix the populations, 15 *elegans* expressing *myo-2p*::*mScarlet* were placed on the bacterial lawn along with 15 of *monodelphis* or *nigoni*. The plates were placed under a fluorescent dissecting scope and allowed to sit for 45 minutes to mitigate any effects of plate movement. The plate was then imaged using white light (for a reference image) and green light (∼546 nm, to capture the *mScarlet* fluorescence). Consistent with previous assays, animals touching two or more individuals were scored as aggregating.

### Orthology analysis

We used the genome sequence and gene annotations from Wormbase^25^ and the *Caenorhabditis* Genome Project to find orthologous gene sets (orthogroups) in non-*elegans Caenorhabditis* species. First, we extracted the longest isoform from every gene using AGAT^26^. Then, using the protein sequence, we constructed orthogroups using OrthoFinder^27,28^ using the sequence aligner, Diamond, in ultra-sensitive mode^29^.

### Statistics and reproducibility

The number for each experiment was based on previous studies and effect size, with each experiment performed with at least 3 independent replicates and each trial performed with matched controls. All data were analyzed and plotted in GraphPad Prism 10 and statistical significance was determined using one-way ANOVA with Tukey’s post-hoc test. For comparisons of two data sets, a two-tailed unpaired t-test was used to compare significance. Error bars on figures represent standard error of the mean (SEM) and p-values are shown in each figure to indicate significance (P<0.05).

## Results

### Social feeding behaviors differ across the *Caenorhabditis* clade

We assayed aggregation and bordering behavior using a quantitative imaging setup^16^. Assays were performed as previously described, with 50 female animals being used instead of 50 hermaphrodites in species that are gonochoristic. In our assay we included four *elegans* control strains; N2 ‘Bristol’ that are solitary, *npr-1(ad609)* that are social feeding, and *npr-1(ad609);nrx-1(wy778)* and CB4856 ‘Hawaiian’ *elegans* that both display intermediate social feeding. To observe any possible trends due to species relatedness, the species were ordered based on their phylogenetic relationship and position. We observed wide variation in mean aggregation behavior amongst members of the genus, with some species like *imperialis* and *virilis* showing relatively high levels of aggregation, although not to the level of the *elegans* social control (*elegans npr-1(ad609)*)(**Figure 2A-B**). Interestingly, *monodelphis* displays similar aggregation as the *elegans* solitary controls (N2) with little to no aggregation (**Figure 2A-B**). Beyond these three examples, we find that the majority of the species have an intermediate level of aggregation, consistent with a modulated aggregation response. Further, we observe a large level of variation between replicates/experiments within some individual species,, namely *japonica, uteleia, and remanei* (**Figure 2A-B**). Somewhat surprisingly, there was no clear pattern between evolutionary relatedness and the observed level of aggregation. In addition to aggregating during social feeding behavior, the *elegans* aggregates tend to be near or in the border of the bacterial lawn where the bacteria is the thickest (**Figure 2A**)^13^. When we examined bordering behavior across the species, we noted two distinct trends. First, a general downward trend in bordering tendency the more distantly related the species are to *elegans*, and second, that the overall trend of widespread variation within some species is also observed for the bordering behavior (**Figure 2A&C**).

To visualize variation among species in both aggregation and bordering phenotypes we generated heat-maps showing the adjusted p-values generated by one-way ANOVA of all combinatorial comparisons (**Figure 3A & B**, respectively). Plotting the phenotypes in a XY plot reveals a relatively broad dispersal, without a clear clustering among closely related species (**Figure 3C**). Overall, the plot emphasizes that while *elegans* N2 and *npr-1*, and *monodelphis* are relative extremes compared to the rest of the species, such extremes occur both in ‘domesticated’ lab and “wild” populations. Consistent with their independent evolution of hermaphroditism^32^, there does not seem to be any significant clustering of the hermaphroditic species. Overall, in characterizing two social feeding behaviors we identified unanticipated levels of variation both among and within species, suggesting our assays may be subject to one or more additional variables that are contributing to variation. Given the wide range of environments from which these species were isolated, we speculated that the lab conditions may represent less than optimal conditions for several species and therefore introduce additional variation.

**Figure 3.**
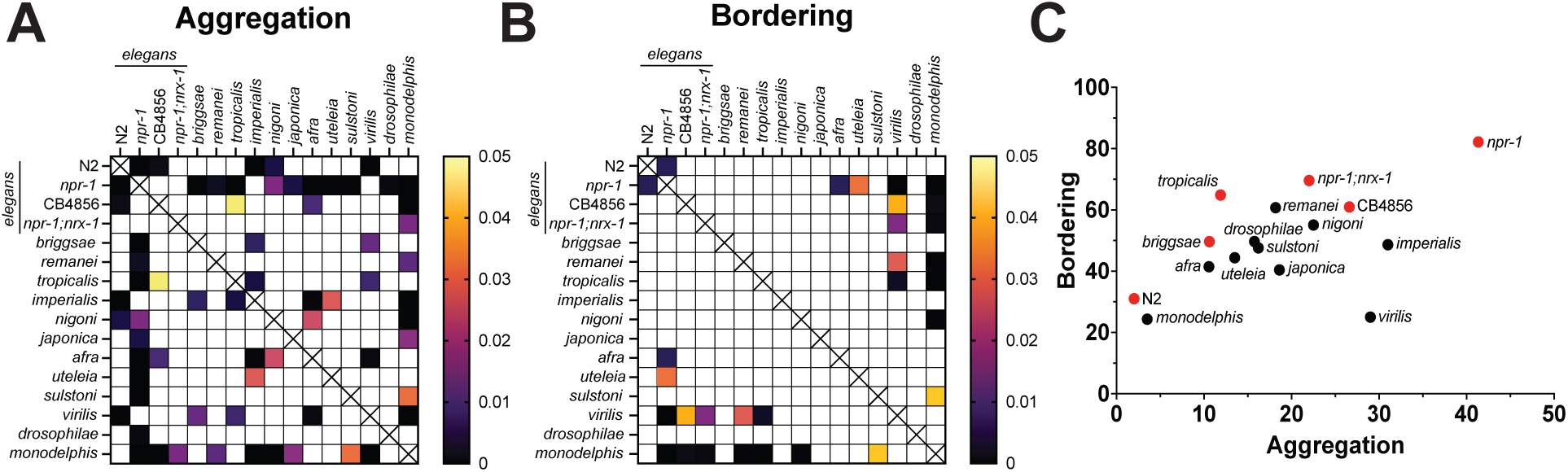
Statistical analysis of phenotypes. **(A)** Heat-map of ANOVA-generated p-values of all pairwise comparisons in aggregation assay. **(B)** Heat-map of ANOVA-generated p-values of all pairwise comparisons in bordering assay. **(C)** XY data plot showing clustering off species when comparing aggregation and bordering phenotypes. As anticipated, *elegans* N2 and *monodelphis*, and *elegans npr-1* represent the low and high extremes of the cluster, respectively. Consistent with the figures above, hermaphroditic species are labeled in red.

### Ecological drivers modulate social feeding behaviors in a species-specific manner

Given the differences in environment and reproductive strategies amongst the species, we reasoned that the makeup of the individual populations could influence social feeding behaviors. To test this hypothesis, we repeated the standard aggregation assays using mixed sex populations (25 males + 25 females, or 25 hermaphrodites in hermaphroditic species with presence of males) (**Figure 4A & B**). For this assay we focused on a subset of the species displaying high level of variation in the female/hermaphrodite only assays, in part to test if a more “ecologically relevant” populations might reduce some of the observed variation. Overall, for most species the mixed sex populations did not alter aggregation behavior compared to the single sex populations (**Figure 4A & B**). However, *monodelphis* and *tropicalis,* showed significantly increased aggregation in the mixed sex population assay compared to the single sex hermaphrodite/female population assay (**Figure 4A**). These results again highlight the impact of local environment and context on variation in behaviors between species, whereas these assays reflect a modulated response to real-time sensory-environment landscape.

**Figure 4.**
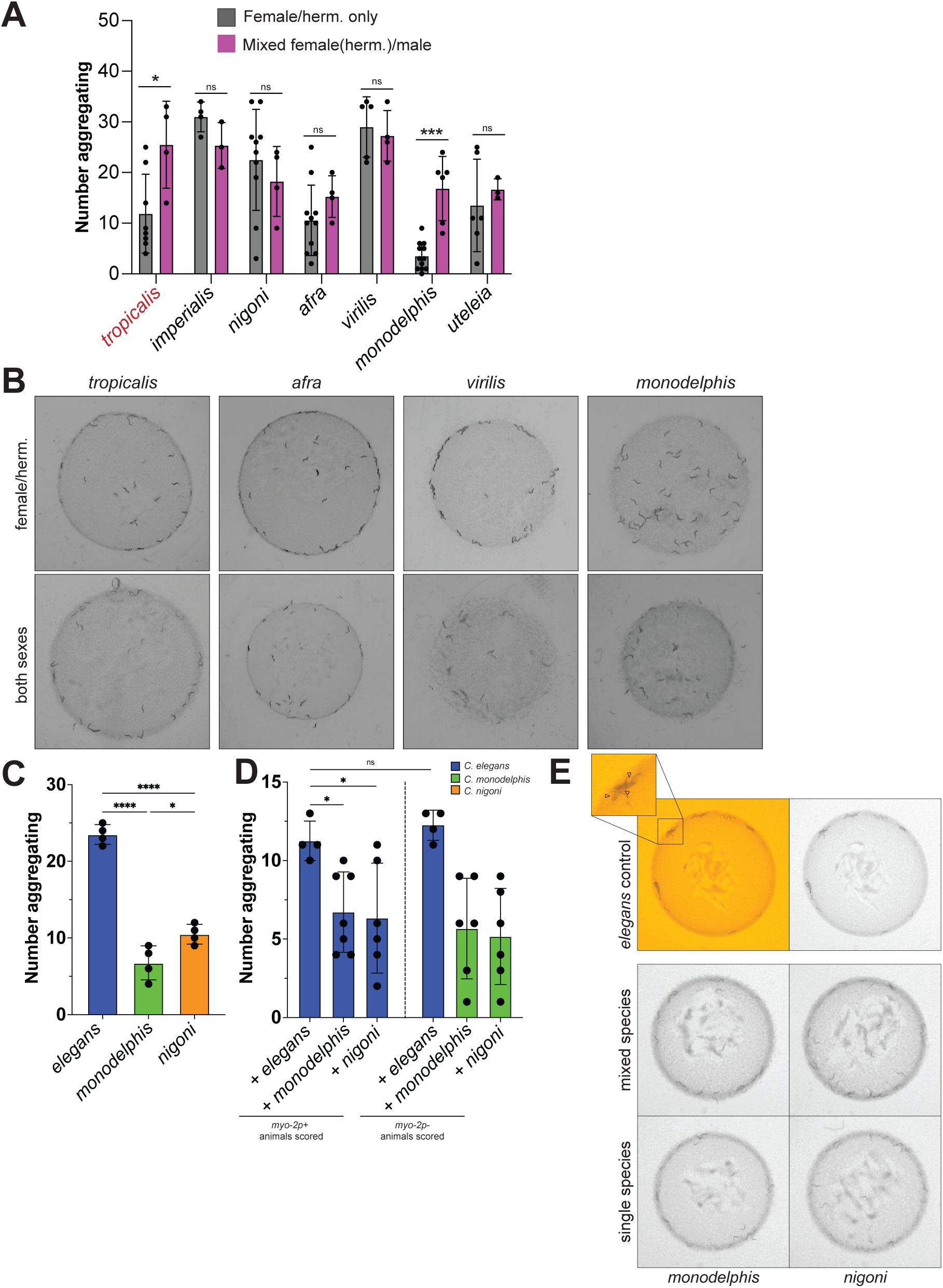
Ecological drivers modulate social feeding behaviors in a species-specific manner. **(A)** Representative images and **(B)** quantification of number of aggregating animals of *Caenorhabditis* species in mixed sex populations. *tropicalis* and *monodelphis* both showed significant differences in aggregation when males were added to the population. Bar graphs represent the mean values across biological replicates (each replicate indicated by a dot). Error bars show stand error of mean. Standard T-test was used for individual comparisons; p-values are indicated in standard format (p=0.05(*), p=0.01(**), p=0.001(***), p= <0.001 (****)). Hermaphroditic species are labeled in red, while gonochoristic species are labeled in black. **(C)** Quantification of 30 animal single-species controls. **D)** Quantification of mixed species assays, where 15 *elegans myo-2p+* animals were mixed with 15 *elegans myo-2p-, monodelphis,* or *nigoni*. Combining *elegans* and *nigoni* significantly lowers aggregation of *myo-2* expressing *elegans* compared to their *elegans* alone control (left), while addition of monodelphis caused an insignificant but trending reduction (left). *monodelphis* and *nigoni* have significantly lower aggregation than their *elegans myo-2p*– control (right). **(E)** Representative images of mixed species assays. *myo-2p* expressing *elegans* are indicated by arrows (top left window).

To explore other ecologically relevant population dynamics (such as interspecies cohabitation and interactions, for example escape from predation^33^), we tested whether combining additional species with *elegans* would influence *elegans* social feeding behavior. We first tested whether a reduced population size (30 animals) or the addition of a *myo-2p* fluorescent transgene would impact the aggregation behaviors of each species. Aggregation in both *nigoni* and *monodelphis* were significantly reduced compared to *elegans* (15 *myo-2p* positive(+) and 15 *myo-2p* negative(-))(**Figure 4C**) and significantly different from one another, which is consistent with the 50 animal aggregation assay results (**Figure 2**). Furthermore, the inclusion of the *myo-2p* transgene did not impact *elegans npr-1*(*ad609*) aggregation behavior. Next, we mixed 15 *myo-2p*+ *elegans npr-1(ad609)* social feeding controls with 15 *myo-2p*-*elegans, monodelphis,* or *nigoni* on a seeded NGM plate and allowed the mixed populations to acclimate for 45 minutes. After the acclimation period, the plates were imaged and assayed for aggregation behavior. We aimed to understand if the mixing of two distinct populations would alter the aggregate feeding dynamics of either or both species. In the mixed species population assays, we noted that addition of *nigoni* or *monodelphis* to the *myo-2p*+ *elegans* population significantly reduced aggregation behavior of *elegans* (**Figure 4D&E)**. Interestingly, we did not observe an impact of *elegans* on behavior of the other species. We reasoned that disruption of *elegans* aggregation behavior could be the result of a number of changes to environmental cues present on the plates that result from mixing populations.

### *Caenorhabditis* clade has high level of conservation in NPR-1 receptor protein

Given the variation in the social feeding behaviors in the *Caenorhabditis* genus, we began to consider what mechanisms might drive species differences. Morphological differences likely drive some of the changes, however, there has been very little characterization in most of the species we assayed. There are several papers examining the development of the vulva ^34–36^, tail morphology^37–39^ and body size^40^, however there is still a significant deficit in the literature. This lack of species characterization is especially apparent when considering the nematode nervous systems, with the only other *Caenorhabditis* species with any significant neuronal characterization being *briggsae*^41,42^. To bypass the lack of circuit-based knowledge within the clade, we leveraged several known genetic drivers of social feeding behavior in *elegans*, notably the neuropeptide receptor NPR-1. To compare *npr-1* across species, we first queried both existing and draft genome assemblies for orthologues using OrthoFinder^28^. Given the size and relative domain conservation of the NPR family of genes in *elegans* we were somewhat surprised to identify a single orthologue of NPR-1 in all species except for *nigoni*, which had multiple related paralogues and was thus excluded from our analysis. To characterize sequence conservation among species we used CLUSTAL V5.0 analysis to perform a multiple sequence alignment to produce an NPR-1 alignment (**Figure 5A**). The alignment showed there is a high level of conservation among the species, even when considering the most evolutionary divergent species, like *monodelphis*. We queried three residues in NPR-1 that when mutated have been shown to induce social behaviors^13^ and found them to be well conserved and wildtype.

**Figure 5.**
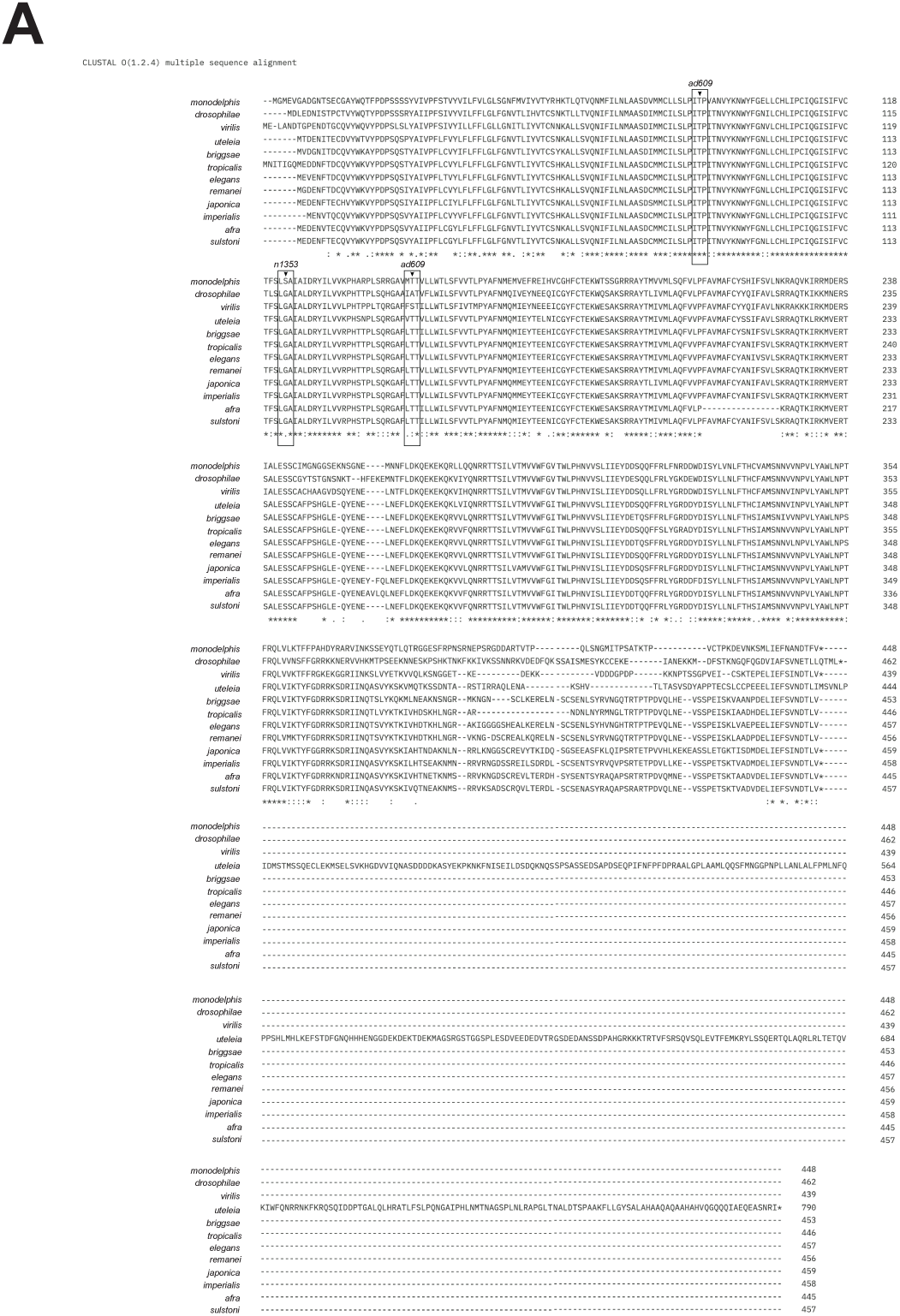
Protein alignment of NPR-1 orthologues in amongst *Caenorhabditis* nematode species. **(A)** NPR-1 orthologues were queried from existing *Caenorhabditis* genomes or assemblies based on sequence similarity. A single orthologue was identified in each species with the exception of *nigoni* which was excluded due to the lack of a confident orthologue. Overall, NPR-1 is relatively well conserved across the clade.

## Discussion

Here, we examined several social foraging behaviors of an evolutionarily diverse clade of *Caenorhabditis* nematodes to better understand how such behaviors differ and may have evolved between species. Across these divergent species of nematodes, we noted several surprising trends within a set of social feeding behaviors that underpin a shared general feeding strategy. Notably, we confirmed *elegans* (*npr-1*) is a relative extreme when compared to several wild isolates and other species. In contrast, most of the species we assayed (including the wild *elegans* isolate CB4856 ‘Hawaiian’ strain) showed intermediate levels of aggregation, while basal taxon *monodelphis* showing little to no aggregation, not dissimilar to *elegans* N2, which is known to be solitary due to a gain of function mutation in the *npr-1* gene. Furthermore, we uncovered a surprising trend when examining bordering behavior among species, notably that the phenotype seems to generally decline the further away the species is in relatedness to *elegans*. While this trend is not perfect, it does again highlight the idea that N2 ‘Bristol’ *elegans* may represent a behavioral outlier when compared with wild *elegans* isolates and most other species.

We noted that moderate to high levels of variation were present in most species when considering both aggregation and bordering phenotypes. The extensive variation present within several of the species raises the possibility that an active mechanism is driving heterogeneity beyond the expected variation present in most behaviors. This unexpected variation could be in part due to genetic heterogeneity present between individuals, maintained in part through selection pressure. *elegans* strains have been artificially selected and lab-evolved (in part because they reproduce clonally), while in comparison, most of the species we assayed represent far more heterogeneous populations. Several species have been backcrossed and inbred extensively from their wild isolates, effectively rendering them lab strains **(Table 1)** after undergoing selection for domestication. We can speculate that some of the species likely retain higher levels of genetic heterogeneity, especially among genetic elements such as single nucleotide variations (SNPs). While the vast majority of SNPs are unlikely to be the causative agent directly underlying a phenotypic change, single nucleotide changes can drastically alter behavior, as highlighted by single nucleotide variation in the neuropeptide receptor NPR-1. Many wild isolates of *elegans* naturally aggregate and display social feeding, but in the lab N2 Bristol strain, a single adenine to guanine substitution in NPR-1 is sufficient to produce a gain of function phenotype, eliminating social behavior and inducing solitary feeding^13^. Lastly, it is not unreasonable to speculate that genetic variants present within individuals may have a modulatory effect within particular behavioral circuits, especially if they are closely associated with critical neuronal genes or regulatory elements^43^. This concept is highlighted in recent work showing mutations in the *daf-19* transcription factor are able to dramatically alter chemosensory and oxygen-sensing circuits leading to a disruption of social feeding behaviors in both *C. elegans* and *P. pacificius*^44^.

Outside of genetic mechanisms, it is very likely that some of the variation seen in social feeding behaviors is a result of the lab environment in which these assays are being performed. In the lab we make every effort to maintain temperature at 20 °C with low to moderate humidity, which is ideal for the growth and maintenance of lab strains like *elegans* and *briggsae*. However, many of the other *Caenorhabditis* species were isolated from drastically different climates and environments and likely underwent adaptation of their environmental-sensory circuits to these divergent conditions in contrast to those in which we assayed them. While testing ecologically relevant temperatures and humidity levels for each species is logistically beyond the scope of this study, one might imagine that social feeding behaviors might be altered if those conditions more accurately reflect what occurs in the wild.

Furthermore, the substrate on which we feed the animals is resoundingly artificial, using a monoculture of a lab strain of *E. coli* (OP50) on an agarose surface is hardly representative of the natural ecology of wild feeding substrates. When considering the natural environments in which these nematodes normally reside, the microbiomes can also be dramatically heterogenous depending on location. Most natural feeding substrates contain a range of bacterial species, some of which are viable food sources and some that may represent nematode pathogens. To reconcile these differences, recent research has characterized food choice preferences in *elegans*, uncovering their preference for *bacillus* species over the traditional lab *E. coli* strains^45^. Even within *E. coli* species, *elegans* tend to prefer nutrient dense strains, likely to supplement for essential nutrients, and potentially leading to changes in metabolism^45,46^. It is not surprising then that different food sources can lead to dramatic variations in behavior. Best studied is the attraction/avoidance response to food, which allows the animals to avoid potentially pathogenic bacteria while optimizing nutrient uptake^47,48^. Lastly, *elegans* has been shown to modulate their feeding behaviors based on the density of food, where both solitary and social strains disperse when food density is strongly diminished^14^, likely as part of a co-dependent circuit detecting both starvation cues and increased oxygen^49^.

The last major factor potentially influencing these behaviors is chemical signaling in the form of environmental secretions and touch stimuli. Although the nature of these chemical modulators is widely unknown, they represent an intriguing mechanism by which individuals can modulate interactions with other animals and their environment. For example, it is well known that several species of nematodes have complex and diverse secretomes, which they use to modulate interactions with their environment. Several studies have shown that parasitic nematodes secrete protein or chemical modulators to downregulate their host’s immune response^50^. Similarly, the *elegans* genome encodes a large number of putative secreted proteins^51^, some of which were isolated via mass spectrometry, identifying a number of novel and nematode specific protein factors of unknown function. In parallel, the chemical secretome of *elegans* revealed a complex biochemical landscape, heavily enriched in putative pheromones, especially ascarosides^52^. Given the unknown function of many of these proteins and chemicals it is not unreasonable to assume at least a subset may be influencing the behaviors of both individuals and populations of individuals. Several ascaroside pheromones for example are well documented to alter population behaviors, modulating hermaphrodite-male attractions and repulsion, mating, and dauer formation in the absence of food^53^. Furthermore, pheromone secretion detected in the environment can also induce the production of EV’s in neuronal tissues of the affected animals, potentially altering behavior^54^. Taken together we must consider that unknown signaling molecules are modulating the interaction between individuals in our social feeding assays. Among the individuals of the same species, these signals may provide a real-time readout of the local sensory environment helping animals to respond and alter their behavior accordingly. Among individuals of different species, the interactions are potentially even more complex. Differences in pheromone usage or chemical secretome might produce novel interactions leading to interesting behavioral variations.

Overall, this work represents the first broad characterization of social feeding behaviors across a large and evolutionarily ancient lineage of nematodes. Given the range of distinct environmental conditions in which these species naturally reside, it may not be surprising that we observed variation among species across several behavioral paradigms. The current circuit mechanism underlying the social feeding behaviors is still trying to fully reconcile how behavior is tuned in response to various sensory cues like food identity/density^14,45,46^ and O_2_/CO_2_ balance^55^. Complicating matters is recent evidence in *P. pacificus* showing that single cues such as elevation (and corresponding depletion of O_2_) can override the established behavioral paradigm and introduce variation^15^. Context-dependent variation is likely to complicate dissection of the genetic and molecular mechanisms of behavioral variation, but presents an exciting opportunity to study real-time plasticity of behaviors and neuronal circuits.

---

**Supplementary Figure 1.**
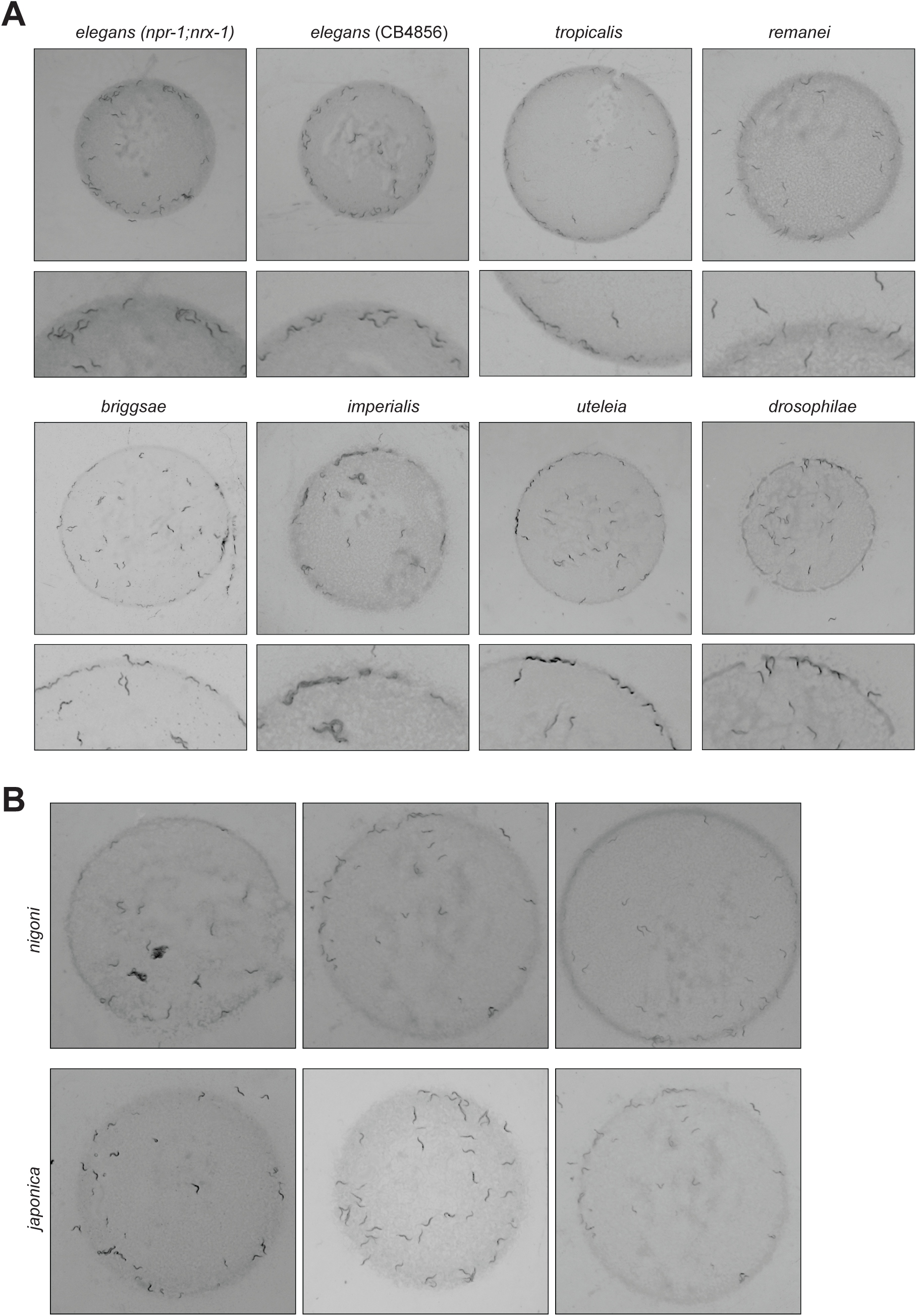
Aggregation behavior is diverse across *Caenorhabditis* clade. **(A)** Representative images of additional species in the *Caenorhabditis* clade. **(B)** Multiple representative images of *nigoni* and *japonica* showing variation in aggregation and bordering behaviors.

## Acknowledgements

The authors thank the labs of John I. Murray, Colin C. Conine, Chris Fang-Yen, David M. Raizen, and Meera V. Sundaram for their feedback on this project. We would also like to thank the Blaxter Lab (The Sanger Institute) for access to the unpublished genomes of several *Caenorhabditis* species. We also thank Yun Ding, John I. Murray, and members of the Hart lab for comments on the manuscript. Some strains were provided by the CGC, which is funded by NIH Office of Research Infrastructure Programs (P40 OD010440). This work was supported in part by NIH R56MH096881 and NIH R35GM146782 (MPH), and the David and Gail Gasser fund (RFHE). CRLL was supported by NSF grant PRFB2305513.

## Author Contributions

DH and MPH conceived and designed the study and experiments, and DH, JP, and RFHE conducted all experiments. DH processed, analyzed, and interpreted all data with help from CRLL. DH wrote the manuscript with assistance from MPH, and all authors reviewed, revised, and approved the manuscript.

